# Octopus arm search strategies over complex surfaces

**DOI:** 10.1101/2023.07.31.551380

**Authors:** Dominic M. Sivitilli, Ariella Zulch, David H Gire

## Abstract

Despite the extreme flexibility of the octopus’s arms and their resulting near infinite possible configurations, the octopus effectively controls its arms during a wide variety of behaviors, including locomotion, foraging, excavation, exploration, and manipulation. If appropriately characterized, the octopus’s biomechanical properties and control strategies could be implemented in the development of a soft robotic limb with the same range of capabilities. When operating without visual feedback, the octopus must rely on the complex chemotactile sensory system within its suckers, and in these conditions sucker recruitment plays a prominent role in search behavior, causing the arm to conform to surface features in the environment. However, how this mechanism is used to search over the complex and convoluted surfaces in the octopus’s natural habitat is unknown. Here, we investigate the strategies the octopus uses to search for a reward hidden among a row of multiple small openings of a task space, and how it uses multiple arms to search three parallel versions of this task space. We found that when the arm encounters multiple openings in a surface, it performs a distal to proximal search pattern, starting with the farthest openings within reach then working its way proximally with a preference for searching each opening in succession. This strategy would allow the octopus to use its highly flexible limbs to perform an exhaustive search pattern over complex surfaces.

## Introduction

The octopus’s arms are remarkably flexible both mechanically and behaviorally. They can bend anywhere along their length in any direction, giving them effectively infinite degrees of freedom (Kennedy et al., 2020; Kier and Smith, 1985; Kier and Stella, 2007), and are employed in virtually all of the octopus’s most common behaviors, including locomotion, foraging, exploration, excavation, and manipulation (Packard et al., 1988; Mather, 1998; Hochner, 2008). The octopus arm therefore presents an exciting biological inspiration for soft robotics. As the biomechanical and computational properties of the arm are revealed, these properties can be implemented in the development of soft robotic limbs with the same range of capabilities.

Suckers stagger along the ventral (bottom-facing) side of each arm typically in the hundreds (Young, 1965; Gutfreund et al., 2006). Most of the octopus’s nervous system exists as a nerve cord extending down the center of each arm (Young, 1963; Rowell, 1966; Budelmann and Young, 1985). Within the nerve cord, localized motor pathways coordinate behaviors activated by the brain while integrating chemotactile information acquired by the suckers. Between the brain and nerve cords, there is a significant reduction in bandwidth. It is estimated that the brain and optical lobes, with around 170 million neurons, communicate with the arms and suckers, with around 350 million neurons, through a pathway of about 172 thousand axons (Young, 1965; Matzner et al., 2000; Rokni and Hochner, 2002; Nesher et al., 2019). Much of the information within the arms and suckers is therefore heavily simplified or lost entirely upon ascending to the brain. Proprioception, as a notable instance, seems to be shared within and between arms, but has no representation within the brain (Wells and Wells, 1957; Wells, 1964; Rowell, 1966). However, the octopus is able to use visual feedback to guide its arm toward a reward without relevant mechanical and chemical cues (Gutnick et al., 2011). With a limited representation of the arrangement of the arms and suckers, the brain is limited in its ability to send a detailed motor plan to the arms. The brain will instead activate arm behavior by signaling a large pool of motor neurons along the nerve cord while relevant sensory input specifies the site of activation (Zullo et al., 2019). Many of these localized motor pathways and their corresponding behaviors, including reaching, recoiling, as well as sucker adhesion, probing, and recruitment, remain largely intact even when the arm is isolated from the brain (Rowell, 1963; Graziadei, 1965; Rowell, 1966; Altman, 1968; Sumbre et al., 2001; Gutfreund et al., 2006; Zullo et al., 2011; Hague et al., 2013; Katz et al., 2021).

The octopus’s arms have been described as highly independent, with arms appearing to execute separate motor patterns in parallel. This has been attributed to the degree of decentralization and autonomy of the arm’s motor pathways (Mather, 1998). While multi-arm behavior and coordination has been a focus of a number of investigations (Buresch et al., 2022; Bidel et al., 2022; Byrne et al., 2006; Levy et al., 2015), few steps have been taken to isolate the arms within separate task spaces to prevent arm interaction and visual occlusion during behavioral recording.

Though sucker recruitment is a behavior that operates in the arms with minimal feedback from the brain, it provides advantages in several contexts. Sucker recruitment occurs when suckers encounter a stimulus, such as a prey item, and neighboring suckers respond by orienting toward this stimulus. These suckers may then recruit their neighbors as more and more suckers serially bend toward the stimulus. This may cause the arm to conform around an object or feature that initiated the recruitment signal (Rowell, 1963; Altman, 1968; Gutfreund et al., 2006; Zullo et al., 2011). Sucker recruitment serves as an effective mechanism for prey capture, for locally evaluating the salience of sensory input, for simplifying arm control (by relying on surface features rather than a central controller to shape the arms), and for searching over surfaces during foraging and exploration (Sivitilli et al., 2023).

**Figure 1.**
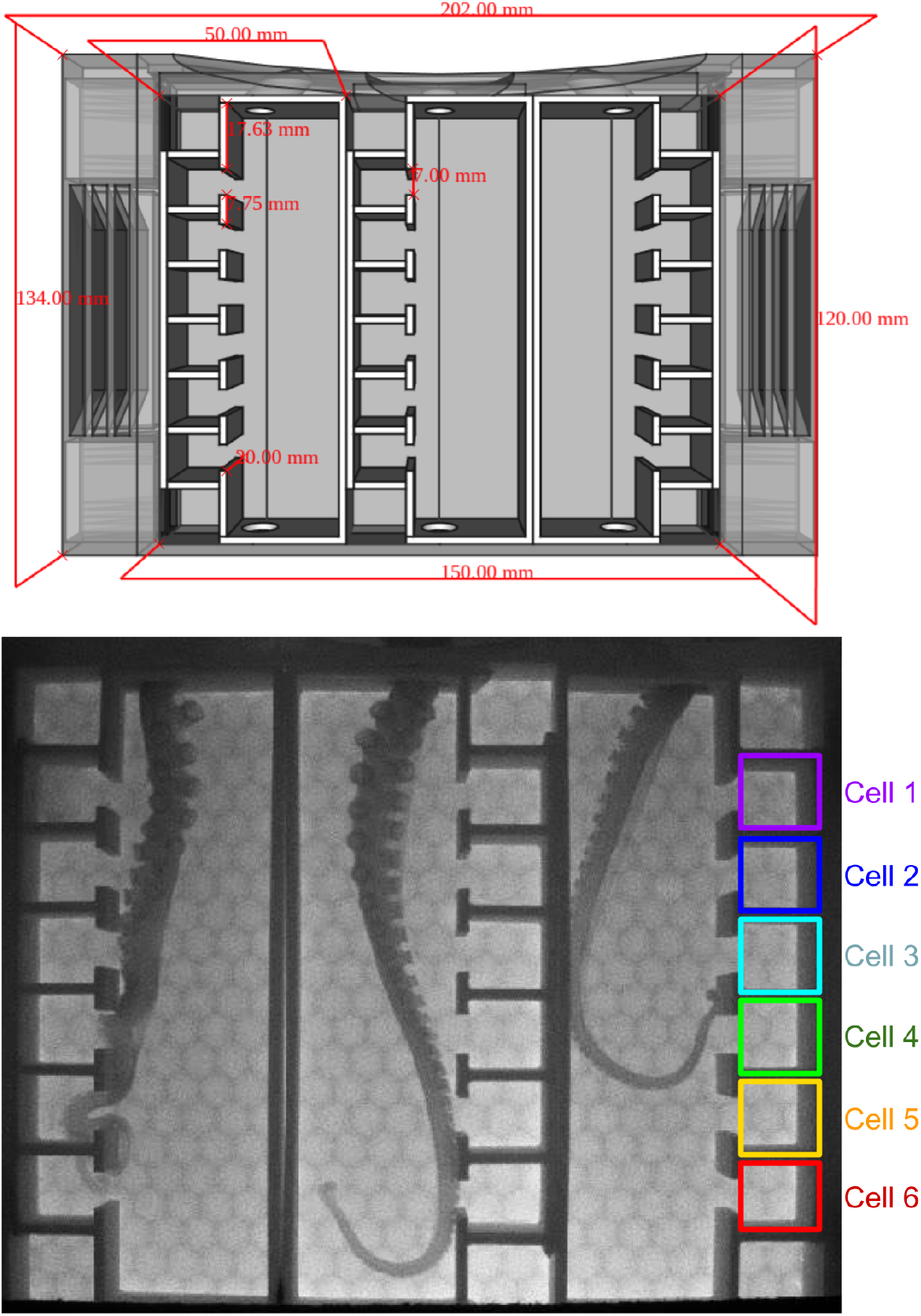
FreeCAD design of the task spaces and frame from a trial recording. Cells were assigned numbers 1-6 based on distance from the entrance.

Beyond the geometrically simple arm tasks used in laboratory settings, exactly how the octopus employs recruitment-led surface conformation when searching over the complex, convoluted surfaces of its natural habitat has not been well-studied. The focus of this investigation was therefore to identify the search patterns employed by the arms within a convoluted task space, which included a row of six regularly spaced openings leading to smaller subsections. Three of these task spaces were arranged in parallel to characterize multiple arms foraging at once. These task spaces were also visually occluded so the octopus could only rely on the chemotactile feedback from its arms and not receive any visual indication of arm shape.

## Methods

Subjects for this study included four male Pacific red octopuses (*Octopus rubescens*) (30g - 115g) and one female giant Pacific octopus (*Enteroctopus dofleini*) (135g), which were collected under an approved scientific collection permit (21-193) through the Washington Department of Fish and Wildlife. Subjects were housed in separate tanks and were transferred to an experimental tank for training and trials. All tanks were supplied with flow-through natural seawater and kept on an artificial lighting schedule that roughly correlated to natural daylight. Subjects were enriched regularly during the course of the experiment. All experiments were carried out in accordance with protocol 4356-02 approved by the University of Washington (UW) Institutional Animal Care and Use Committee (IACUC).

The task was designed in FreeCAD and 3D printed with white polylactic acid (PLA) filament. The task was secured against the side of the tank using magnets so the interior task space was visible to a camera outside the tank but not to the octopus. The task interior was shaped as an empty narrow box with three parallel circular entrances at the top. Different versions of the task space could therefore be inserted into the interior.

Subjects were trained to reach into the task by leaving food on the top of the task. Once they readily approached and reached into the task, a training task space was placed inside. This was small enough for the reward to be easily found within reach. Once the subjects readily approached and reached into the task, the training task space was replaced by the three experimental task spaces and trials began. Subjects were given five minutes to search the task while the task space was recorded, plus an additional minute following successfully finding a reward. Each subject was given as many of these opportunities as they needed to find at least one reward hidden within the task space 12 times without consuming more than 2% of their mass in food daily.

Each of the three task spaces contained a large rectangular section with six evenly spaced openings leading to separate square subsections (cells) aligned in a row on one side. Rewards were assigned to a combination of the three task spaces and cells based on a pseudorandom pattern. No more than one reward was hidden in each task space, and if multiple task spaces contained a reward, the rewards were hidden within the same cell number. For the first six trials, two to three of the task spaces contained a reward and for the last six trials, only one task space contained a reward. The task spaces were oriented such that cells for the left task space were on the left side of the camera’s view, cells for the right task space were on the right side, and the middle alternated in its orientation.

Arm behavior during trials was recorded with a CMOS camera (Imaging Source, model DMK 37BUX287) at 250 FPS, and videos were downsampled to 25 FPS for analysis. The arms were tracked using background subtraction and task cells were segmented both with automated Python routines. A number of metrics were used to characterize search patterns. Arm size was calculated as the number of pixels of the arm’s two dimensional shape. Arm reach was calculated as the farthest distance into the task space occupied by the arm. Arm curvature was defined as the Kolmogorov-Smirnov (K-S) statistic comparing the distribution of the skeletonized arm to a straight line projecting away from the task space entrance, with each set of pixel values representing the distance of each pixel from the task space entrance (based on a euclidean distance transform relative to the entrance). Surface conformation was defined as the amount of overlap between the arm and a single pixel-width outline of the task space as a proportion of the arm’s size. Cell occupancy was calculated as a binary value representing any overlap between the arm’s shape and the individual cells.

## Results

To broadly characterize the arms’ search pattern, a time lagged cross correlation related arm reach, arm curvature, surface conformation, and cell occupancy for all trials and subjects. For each trial this generated a series of distributions of Pearson’s correlation coefficient reflecting the synchrony of arm reach and the other metrics over a range of lagged time intervals. These distributions were then averaged across trials producing Figure 2a. This shows a general profile of the arms’ search pattern for 20 seconds leading up to and following occurrences of arm reach. Reaching is shown to be closely aligned with distal cell occupancy and followed by proximal cell occupancy, with more proximal cells having a longer occupancy time. Curvature and surface conformation lag behind reaching along with proximal cell occupancy, showing that while distal cells are occupied with an outstretched arm, the arm is not conforming to the shape of the cells in this state. As the arm progresses proximally, it shows greater curvature and greater degree of conformation to cells.

**Figure 2.**
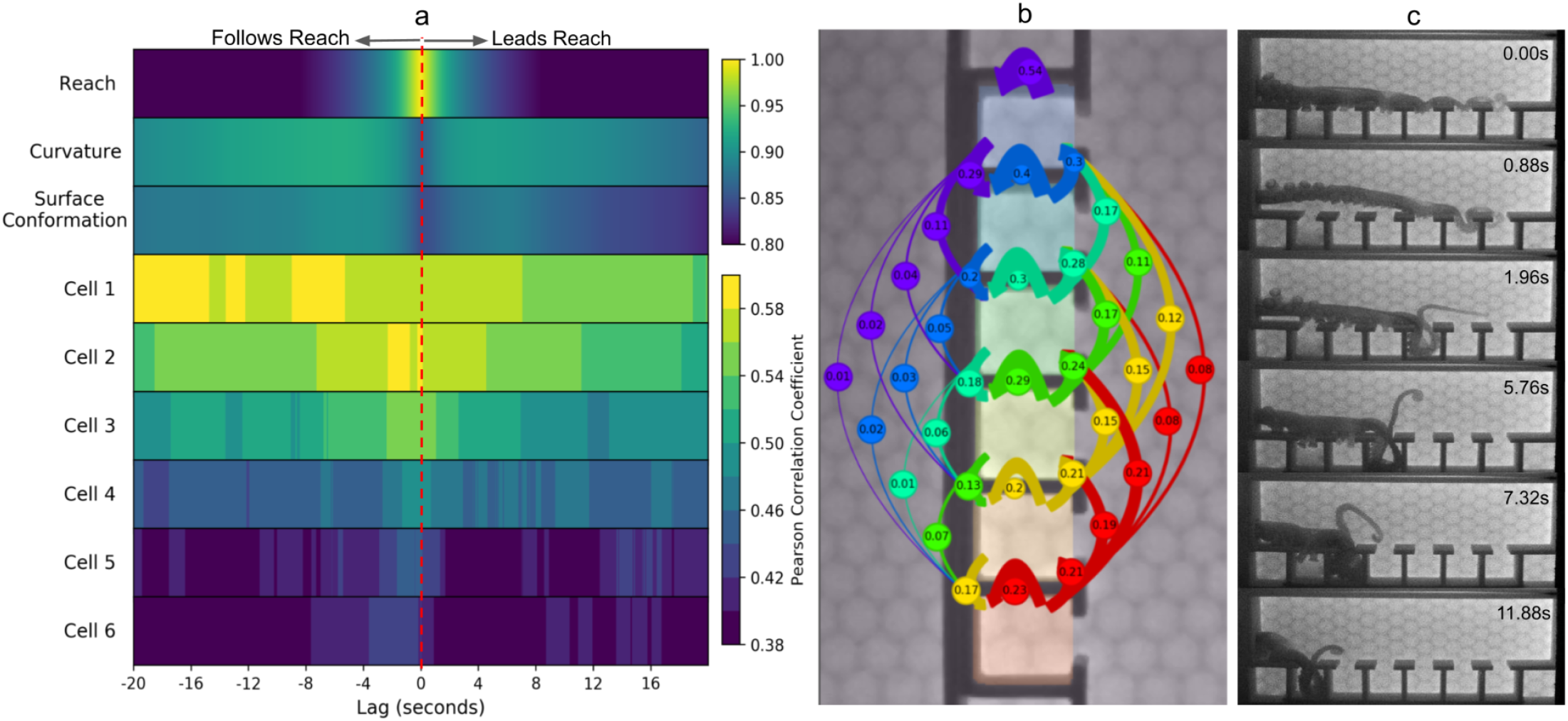
(a) Average time lagged cross-correlation of all trials showing occurrences of arm curvature, arm surface conformation, and cell occupancy and how they correlate with occurrences of arm reaching (at Lag = 0) 20 seconds ahead of reaching (right of plot) and 20 seconds following reaching (left of plot). (b) Transition probability diagram showing the probability of the arm’s most recently occupied cell given its previously occupied cell. (c) An arm performing an exhaustive search pattern while a food reward was hidden in cell 1.

To characterize patterns of cell occupancy, sequences were generated based on the most recent cell occupied while the arms searched the task space. From these sequences a transition matrix was created, which is shown diagrammatically in Figure 2b. In general, between moving proximally, moving distally, and staying in place, the arms were most likely to move proximally, and the most likely transition during proximal movement was to the immediate proximal cell.

To assess the degree of arm independence, the Pearson correlation coefficient was calculated for arm size, reach, curvature, and surface conformation from all occurrences of arms foraging simultaneously in separate task spaces (“parallel”), and for all occurrences of arms foraging separately (“separate”). From these values, two distributions were created for each metric, one representing the correlation profile of parallel arms and other representing the correlation profile of separate arms. A K-S test was then used to compare the two distributions for each metric. Results are shown in Figure 3. Distributions of arm reach, curvature, and surface conformation showed no significant difference, while the distribution of arm size showed that parallel arms were significantly more likely to be anticorrelated (K-S, p = 0.003). Overall these results suggest that for reach, curvature and surface conformation, arms searching in parallel are not any more likely to be correlated or anticorrelated than arms foraging separately, while it is more likely for parallel arms to alternate in the how much of each arm is searching their respective task space.

**Figure 3.**
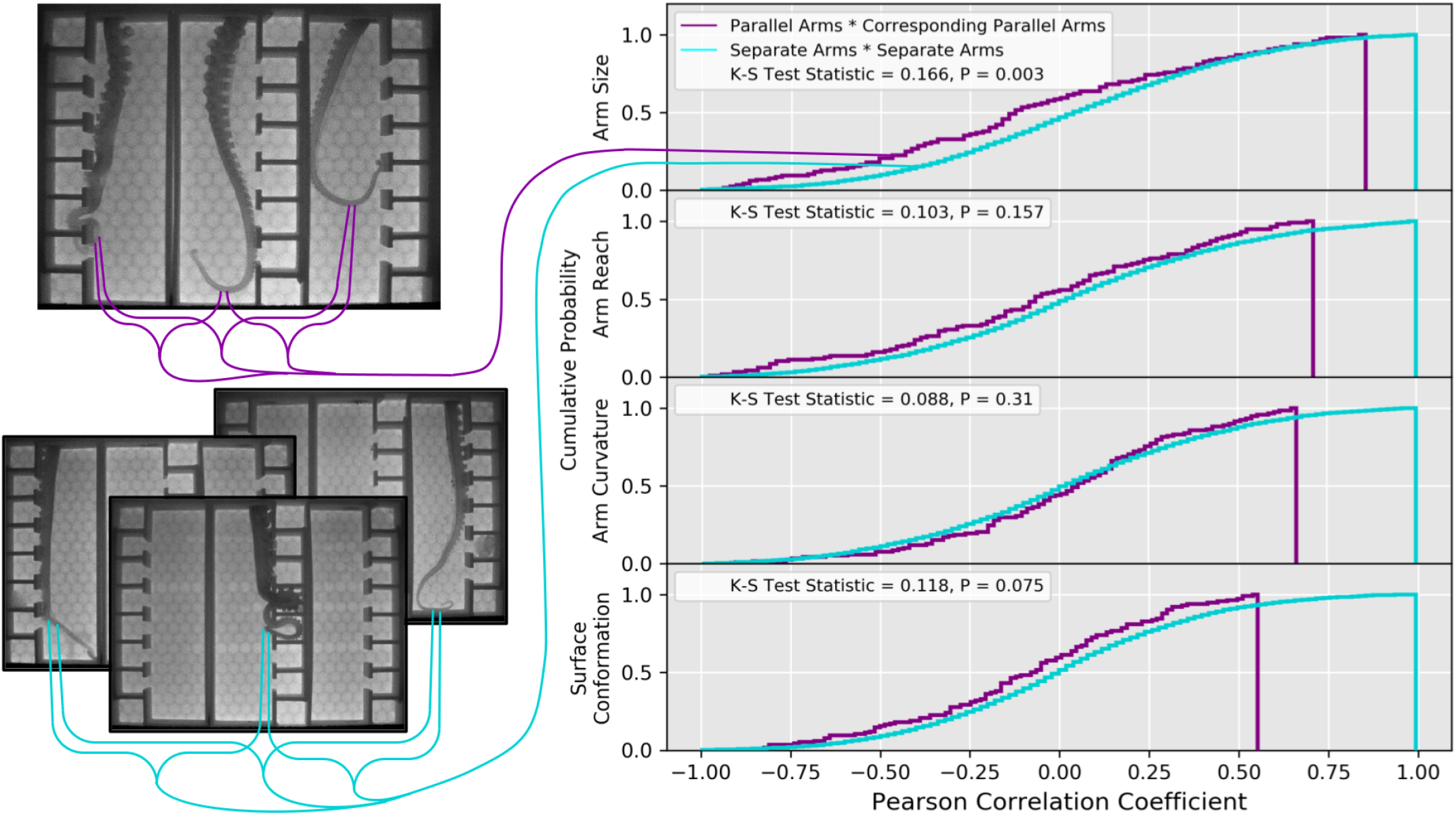
Cumulative probability distributions of Pearson correlation coefficients comparing arms foraging in parallel with their corresponding parallel arm and comparing arms foraging separately. These distributions were compared with a K-S test. Arm size among parallel arms was significantly more likely to be anticorrelated (p = 0.003), while no significant difference emerged among the other metrics.

## Discussion

While recruitment-led search patterns over simple surfaces would seem to follow a predictable pattern of conforming to the surface shape with a particular preference for concave edges (Sivitilli et al., 2023), in its natural environment the octopus would encounter a variety of complex surfaces, including crevices of various shapes, sizes and regularities. Hidden among these surfaces are prey which the octopus is able to locate and extract using just its arms and suckers. How does the octopus adapt its search strategy to successfully forage over these surfaces? Our results reveal a possible strategy.

The three task spaces in this study included a row of evenly spaced openings, and a food reward was hidden within one of these openings. Complete conformation of the arm to the task space surface, an effective strategy when searching over simpler geometries, would have been unlikely to lead to a successful trial if the arm was not long enough to follow the outline of the task to the reward location. This is particularly true for rewards hidden in distal cells. Instead, it was common for the arm to extend across the whole surface, then for the suckers to probe through the openings. The arm would then search in a single cell or multiple cells at a time, and this appeared to result from sucker recruitment-led conformation causing the arm to follow the surface into the opening. These recruitment events occurred within isolated sections of the arm, suggesting that when encountering multiple openings in a surface, the arm can isolate recruitment to specific sections to explore into a single opening rather than needing to continuously conform to the entire surface at once.

The results from the time lagged cross correlation and transition matrix revealed a general search strategy including the order that cells were occupied. When extended, the arm initially explored distal openings then worked its way proximally as it showed progressively greater curvature and surface conformation. Optimizing surface conformation when searching proximal openings makes sense as using this strategy for openings easily within reach would not limit the arm’s ability to reach more distal openings. While cells were commonly skipped, particularly in the proximal direction, the most likely transition for cells 2-5 apart from not moving was to the neighboring proximal cell. Figure 2c shows an example of an arm systematically searching through cells 5-1.

From these results we propose the following mechanism. When outstretched along a surface, the suckers encounter and signal the location of irregularities, which in this case were openings in the task space surface. A mechanism then prioritizes these openings, then an inhibitory signal is sent to the arm section distal to the opening of highest priority while localized sucker recruitment conforms the now free distal arm section into the opening. In the case of equally salient openings, we predict that priority defaults to the distal-most opening. Once the opening is explored, this process is repeated for the next opening and so on. This allows the arm to perform an exhaustive search of all encountered openings by simply alternating between a distally oriented inhibitory signal and sucker recruitment.

Realistically, salience would never be truly equal at multiple points along the arm. Chemical cues, fluid motion, and the size and configuration of the arm and suckers are all likely factored into salience. This is likely the source of cells being skipped or taking immediate precedence. Interestingly, the arm commonly performed an exhaustive search pattern even with the likely presence of an attractive chemical indicating the presence of food within a proximal cell (see Figure 2c). If the suckers are relaying the presence of food, this behavior may suggest that the octopus chooses to perform the entire search pattern rather than prioritizing reward cells, possibly because these cells already fall along an intended search path. This behavior further suggests the presence of an exploration-exploitation balance that may be affected by satiation levels and predation threat.

The narrow architecture of the present task space limits lateral motion in the arm, so while the search mechanism proposed here would be effective in narrow spaces, it fails to account for lateral search over wider surfaces. However, results reported by Sivitilli et al. (2023) show a preference for the arm to search along concave edges. These combined results therefore suggest that the arm preferentially searches along concave surface contours, then proceeds to search through smaller openings along the contour using the mechanism described here.

The degree of arm independence during the task was assessed by creating a distribution of Pearson correlation coefficients comparing arm size, reach, curvature, and surface conformation between arms foraging in parallel and between arms foraging separately, and comparing these distributions using a K-S test. Our results show that reach, curvature and surface conformation among arms foraging in parallel are not significantly more or less correlated than arms foraging separately. However, parallel arms are more likely to be anticorrelated in size compared to separate arms. This likely reflects a feature of the task itself. The closer the octopus is to any one of the entrances, the more of its arm it can reach into the corresponding task space, but this limits the size of the arm it can reach into the others. Unfortunately, arms cannot be easily identified using the methods employed in this study, so we cannot assess whether varying patterns emerge when comparing neighboring arms foraging in parallel to non-neighboring arms foraging in parallel.

Our findings highlight a behavioral advantage of limitless degrees of freedom. By employing a simple behavioral program in its arms combining sucker recruitment with distal inhibition, the octopus can perform an exhaustive search pattern over complex surfaces.

## Acknowledgements

We would like to thank Dr. Kirt Onthank, the Onthank Lab, Willem Weertman, Joey Ullmann, and Julia Kobelt for their valuable help with animal care and collections.

